# Network topology of neural systems supporting avalanche dynamics predicts stimulus propagation and recovery

**DOI:** 10.1101/504761

**Authors:** Harang Ju, Jason Z. Kim, Danielle S. Bassett

## Abstract

Many neural systems display avalanche behavior characterized by uninterrupted sequences of neuronal firing whose distributions of size and durations are heavy-tailed. Theoretical models of such systems suggest that these dynamics support optimal information transmission and storage. However, the unknown role of network structure precludes an understanding of how variations in network topology manifest in neural dynamics and either support or impinge upon information processing. Here, using a generalized spiking model, we develop a mechanistic understanding of how network topology supports information processing through network dynamics. First, we show how network topology determines network dynamics by analytically and numerically demonstrating that network topology can be designed to propagate stimulus patterns for long durations. We then identify strongly connected cycles as empirically observable network motifs that are prevalent in such networks. Next, we show that within a network, mathematical intuitions from network control theory are tightly linked with dynamics initiated by node-specific stimulation and can identify stimuli that promote long-lasting cascades. Finally, we use these network-based metrics and control-based stimuli to demonstrate that long-lasting cascade dynamics facilitate delayed recovery of stimulus patterns from network activity, as measured by mutual information. Collectively, our results provide evidence that cortical networks are structured with architectural motifs that support long-lasting propagation and recovery of a few crucial patterns of stimulation, especially those consisting of activity in highly controllable neurons. Broadly, our results imply that avalanching neural networks could contribute to cognitive faculties that require persistent activation of neuronal patterns, such as working memory or attention.

## Introduction

A central question in neuroscience is how connections between neurons determine patterns of neurophysiological activity that support organism function. Networks of neurons receive incoming stimuli and perform computations to shape cognition and behavior, such as the visual recognition of faces in regulating social behavior [1]. While many studies laud the ultimate goal of determining how brain network topology supports information processing [29,77], it remains challenging to empirically study the direct interactions between neural dynamics, connectivity, and computation. Indeed, neural connections and their underlying computational function have often been inferred through neural dynamics, and formal studies probing mechanistic relations among the three components have remained largely theoretical [8,30,74].

A characteristic empirical feature of many neural systems is avalanche dynamics. Avalanches comprise cascades of neural activity, in which groups of spikes occur uninterrupted in time. These cascades often display heavy-tailed distributions of *size*, as given by the number of neurons that spike in a cascade, and of *duration*, as given by the period of continuous activity [6]. Such distributions, commonly reported to be power-law like, have been detected using a range of methods and have been observed *in vitro* [6,23], *in vivo* [7,18,49,51,59], and *ex vivo* [56] in a variety of organisms, including humans. In a complementary line of theoretical work, power-law distributions in size and duration have been linked to self-organized criticality [4], and systems of this type are said to have optimal information storage [24], optimal information transmission [6], optimal dynamic range [36,57], and computational power [9].

While neural dynamics can be easily characterized, often left implicit in the analysis of neuronal networks are (i) the computational function of avalanches without the assumption of self-organized criticality and (ii) the network structure that supports avalanche dynamics. The dynamics of long cascades characteristic of avalanches have yet to be connected directly to information processing. Instead, they are commonly treated as evidence for the criticality of avalanching neural networks, although whether self-organized criticality is relevant for the brain is debated [16,52,71,72]. Moreover, empirical evidence of strongly and bidirectionally connected pairs of neurons [37,38,75] and higher order network motifs in local and macroscale circuits [42, 48, 60, 63] has yet to be linked to the avalanching phenomenon that is purportedly supported by those same circuits. Although important work addresses how a balance between excitation and inhibition is required for avalanching dynamics in rat slices [40], cortical cell cultures during development [69], and computational models [50], the actual network topology of systems that support avalanches remains largely unexplored. In summary, it is not yet known which network topologies might produce avalanching dynamics, or how those dynamics could lead to certain types of information processing.

Here, we address this gap in knowledge through a series of analyses and numerical simulations of a stochastic model [6,24,25,52] instantiated upon various network topologies. First, as a holistic theoretical framework, we analyze activity propagation in the model via state transitions in a Markov chain to show that sustained activity is required for a network to maintain information about stimuli over time. We then present an analytical constraint of cascade duration by network topology. A precise assessment of this relation in large neural systems is not numerically feasible, and thus we develop further intuitions from estimation approaches. Given this topological constraint of neural dynamics, we also find that cycles and strong connections in cycles – both of which are empirically observed network motifs – are notable contributors to long cascade duration. Finally, we use simulations on 4 commonly studied generative graph models to probe the relations among network topology, cascade duration, and the information about stimuli that is maintained in a network. Using mutual information to measure information maintenance, we show that long cascade durations allow recovery of stimuli via persistent network activity. Collectively, our findings show that the network topology reported extensively in the empirical literature can produce complex, avalanching dynamics in addition to the well-known attractor dynamics [3,11,30]. Moreover, these dynamics support the persistent activation of a cluster of neurons, which in turn allows for the discrimination of stimulus patterns implicated in working memory [14,15,20].

## Mathematical Framework

We begin with the stipulation of a network as well as a dynamical process that occurs atop the network. We formalize the notion of a network as a directed graph 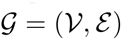 in which neurons are represented as nodes 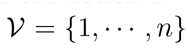 and neuron-to-neuron connections are represented as edges 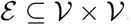. The weighted and directed adjacency matrix *A* = [*a_ij_*] thus encodes the edge weights from neuron *j* to neuron *i*. To model neuronal cascades, we next stipulate a stochastic version of the McCulloch-Pitts neuron. In the McCulloch-Pitts model, a neuron receives inputs scaled by the weights of the edges, whose sum is normalized to 1, and sums the scaled inputs to produce an output via an activation function. Here, the activation function is a Bernoulli process, where probability *p* is the sum of the scaled inputs. The model is similar to the branching model that has been applied to networks to study neuronal avalanches, but limits the maximum firing at one time step to 1 to ensure greater biological realism [6,24,25,52]. The state of an *n-*neuron network is a binary vector ***y**(t)* ∈ {0, 1}*^n^* such that each element indicates whether a neuron fired at time *t* and evolves as

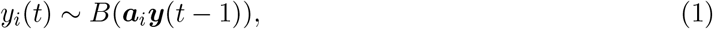

where *B(r)* is a Bernoulli process with probability *r*, and ***a**_i_* is the *i^th^* row vector of *A.*

The model that we consider can be represented as a Markov chain with states ***s**^i^* ∈ {0, 1}*^n^* representing all possible patterns of spikes in the network, and with state ***s***^1^ = 0 representing the zero state. The column vector 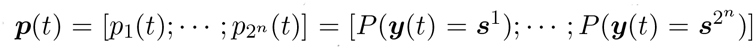 represents the probability that the network exists in any state ***s**^i^* at time *t*. The transition matrix *T* governs

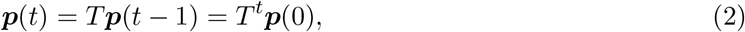

where each entry 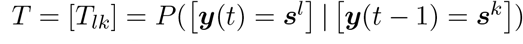 represents the transition probability from state *k* to state *l*. See Methods for details regarding the computation of the matrix *T*.

The Markov representation above makes explicit the relationship between the network *A* and the stimulus propagation and discrimination. The process stated in Equation 1 determines a unique map from adjacency matrix *A* to transition matrix *T*. Given an initial distribution of states, i.e., the stimulus patterns, 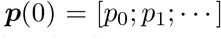, the fraction of neuronal avalanches that terminate by time *t* is simply given by the first entry of ***p***(*t*) = *T^t^**p***(0). Similarly, the discrimination between network states propagated from stimulus ***y***(0) = ***s**^i^* and from ***y***(0) = ***s**^j^* from some measurement ***y***(*t*) depends upon the similarity between probability vectors ***p**_i_*(*t*) = *T^t^**s**^i^* and ***p**_j_*(*t*) = *T^t^**s**^j^*. For quickly decaying systems, ***p**_i_*(*t*) and ***p**_j_*(*t*) will both have a high probability of being in the zero state *s*^1^, inherently reducing discriminability. Hence, the architecture of the network *A* constrains the amount of persisting activity that permits discrimination of the initial spiking distribution ***p***(0).

Because computing a Markov chain is intractable for large network sizes, we instead take advantage of the linear relation in the process stated in Equation 1 to estimate stochastic behavior.

Specifically, the average activity generated by the stochastic model can be written as 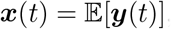, and it is straightforward to show that this average network state obeys

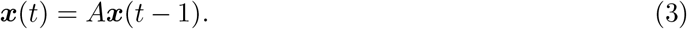

(see Methods for a formal proof). Equation 3 offers a natural intuition: the average behavior of the stochastic branching model follows linear dynamics. Such a relationship allows the application of rich mathematical principles of linear dynamical systems to describe average stochastic dynamics of the model, especially in the context of neuronal avalanches.

To illustrate the linear relation, we perform numerical simulations of the model, and we compare simulated cascades to the average network state estimated by a linear dynamical system. In both cases, we instantiate the dynamics on a weighted random network comprised of 10 nodes (Figure 1) [17]. The important network parameters for all simulations are listed in the Supplemental Materials. We simulate the branching model dynamics 1,000 times over 15 time steps starting with the same initial condition *y*(0) (Figure 1b). Note that we can also consider this initial condition to be the stimulus. We average the activity at each node and time step ***y**_j_(t)* across simulations to generate a numerical estimate of the time-evolution of the average network state (Figure 1c). Then using the linear dynamical system starting with the same initial condition ***x***(0) = ***y***(0), we calculated the number of spikes per neuron per time step as a second estimate of the average network state (Figure 1d). We find that the difference between the states predicted by the simulation of the branching model and the states predicted by the linear system is consistently small across a range of network sizes for fixed density (Figure 1e-f). These results illustrate the accuracy of the linear estimation of the dynamics of the stochastic model.

**Figure 1.**
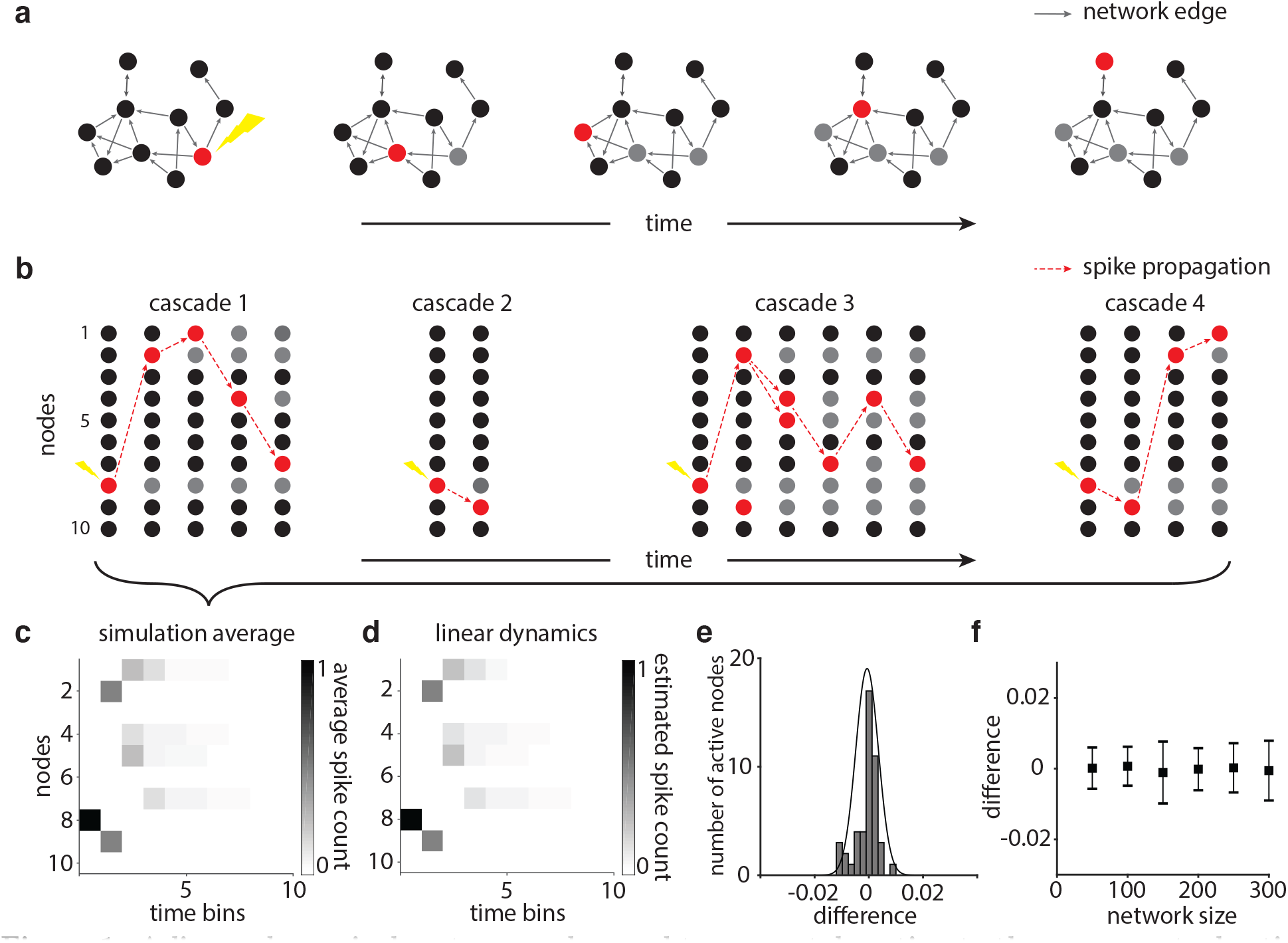
A linear dynamical system can be used to accurately estimate the average stochastic spiking of neurons in a branching process model. **a**, An example schematic of the cascade generated by an external stimulus shown in the leftmost portion of panel **b.** The red nodes indicate active nodes, and the gray nodes indicate nodes that have already been active. **b**, Examples of different cascades generated by stimulating the same node of a fixed weighted random network. In this case, we stimulate node 8 and note that therefore node 8 is the only active node in the first time bin, but thereafter the cascades may differ. **c**, The activity of each node and time step, averaged over 1,000 simulations on the same weighted random network and with the same stimulation node used to produce the results shown in panel **b**. **d**, The linear dynamical system with the same weighted random network produces an estimation for spike counts of the stochastic branching model. **e**, A histogram of the difference between the cascades produced by the simulation average shown in panel **c** and the spike counts stipulated by the linear dynamical system shown in panel **d**. The normal fit has a mean of −6.4 × 10^-4^ and a standard deviation of 4.2 × 10^-3^. **f**, For weighted random networks of size 50, 100, 150, 200, 250, and 300 nodes, all with fractional connectivity of 0.2, we show the differences between the average cascade activity and the estimation via linear dynamics. The error bars indicate standard deviations, and the means are 2.094 × 10^-4^ (50 nodes), −6.76 × 10^-5^ (100 nodes), −9.7 × 10^-6^ (150 nodes), −8.0 × 10^-5^, 5.75 × 10^-5^ (200 nodes), 5.75 × 10^-5^ (250 nodes), and −2.05 × 10^-5^ (300 nodes).

## Results

### Network topology constrains cascade duration

As the first step towards uncovering relations between architecture and avalanche dynamics, we provide an accurate relationship between network topology and cascade duration using intuitions grounded in the theory of linear dynamical systems. To analytically describe cascade dynamics, we take advantage of the linear relation in Equation 3. Because a cascade is alive if any neuron is spiking, the probability that a cascade is alive at time *t* is given by

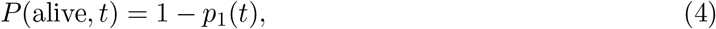

where *p*_1_(*t*) is the first element of the state probability vector ***p***(*t*) = *T^t^**p***(0) from Equation 2. Given a stimulus pattern ***y***(0) and a large number of trials, the fraction of cascades alive at time *t* approaches *P*(alive,*t*) (Figure 2a). Because the transition matrix *T* is derived from *A*, the propagation of the initial stimulus ***y***(0) is determined by the network *A*.

**Figure 2.**
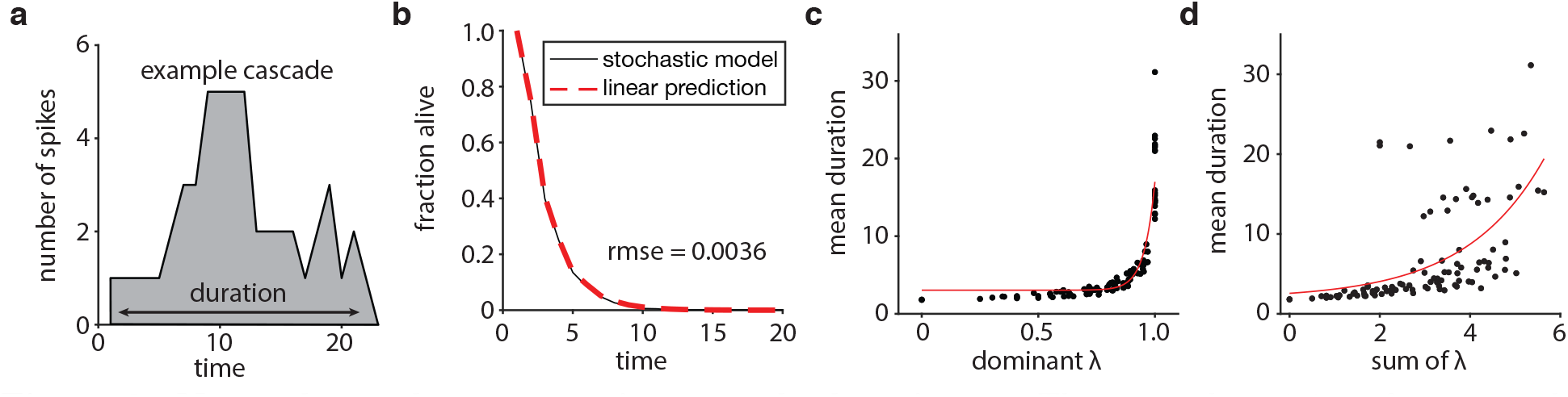
Network topology constrains cascade duration. **a**, The network activity of an example cascade on a weighted random network. The cascade duration is defined as the number of time steps *t* between the point at which the first spike occurs after a time step of quiescence, and the point at which the last spike occurs, followed by a time step of quiescence. **b**, The dynamics of the stochastic model can be accurately predicted by linear estimation. The plot shows the fraction of cascades alive at time *t* from 1,000 trials of stimulating a node in a single instantiation of a 10-node weighted random graph. The root-mean-square error between the stochastic simulation and the linear prediction is 0.0036. **c**, Networks with larger dominant eigenvalues allow longer cascades. The plot shows the mean duration of cascades as a function of the dominant eigenvalue of the network. Each of the 100 points reflects a single weighted random network composed of 10 nodes. The value of each point is the mean across the 1,000 values obtained from stimulating each of 1,000 randomly selected nodes from the graph. Note that because the network is composed of 10 nodes, and 1,000 selections are made, each node is selected multiple times. **d**, Networks with larger sum of eigenvalues allow longer cascades. The plot shows the mean duration of cascades as a function of the sum of eigenvalues of the network. Similar to panel c, each of the 100 points reflects a single weighted random network composed of 10 nodes. The value of each point is the mean across the 1,000 values obtained from stimulating each of 1,000 randomly selected nodes from the graph.

To numerically assess *P*(alive, *t*), we simulated the 10-node, weighted random network used in Figure 1 and observed little difference between the stochastic and predicted dynamics. For each of 1,000 trials, we stimulated single neurons at *t* = 1, and at each time step (from a maximum of 100), we calculated the fraction of cascades alive and *P*(alive, *t*). We found that the root-mean-square error (RMSE) between the linear prediction and the stochastic model was 0.0036 (Figure 2b). To determine the generalizability of our observations, we extended this analysis to an ensemble of 120 networks, separated into 30 instantiations of four different graph topologies chosen for their relevance to neuronal architectures: a weighted random graph, a ring lattice graph, a modular graph with 4 communities, and a Watts-Strogatz graph (see Methods). For the four graph topologies, we observed that the average RMSEs were less than 8.5 × 10^-3^. Taken together, these results indicate a tight link between network structure *A* and cascade duration derived from the network dynamics *T*.

To more generally describe cascade behavior given just the topology *A*, we can decompose the weight matrix *A* into eigenvalues and eigenvectors to identify the elementary modes of activity propagation. One metric we can use to identify the maximum constraint on the dominant propagation of activity is the dominant eigenvalue ⋋_1_, where

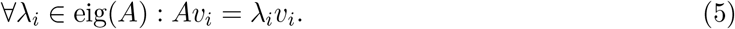

The dominant eigenvalue ⋋_1_ scales the dominant eigenvector *υ*_1_, which constrains the most persistent mode, or vector, of activity propagation in the network. In addition, other modes of activity propagation can affect cascade dynamics. Thus, for a more holistic view of activity propagation in the network, we can compute the sum of the absolute value of all eigenvalues, |**⋋**|_*i*_. This sum represents the rate of decay of all elementary modes of activity propagation. We present the dominant eigenvalue and the sum of all eigenvalues as two metrics of estimating activity propagation in a network.

To numerically demonstrate the utility of the two metrics in explaining cascade duration, we simulated 100 networks with 10 nodes and weighted random graph topology. For each network, we stimulated individual neurons at random, and the resultant cascade was allowed to propagate for 100 time steps. Because the graphs are randomly constructed, they had a range of dominant eigenvalues from 0 to 1.0 (Figure 2c). The mean duration of cascades is approximately linearly correlated on a log scale with the dominant eigenvalue and with the sum of eigenvalues (Pearson’s correlation coefficients *r =* 0.7830 and *r =* 0.7073, respectively; Figure 2c,d). Notably, these relations can inform how one would tune the network *A* to produce cascades of long duration.

### Network motifs: cycles and connection strength

Having demonstrated in the previous section that cascade duration can be predicted from the network topology, we next turn to a deeper examination of which specific features of a network’s topology and geometry can support a heavy-tailed distribution of cascade duration. Note that we use the phrase *network topology* to indicate the arrangement of binary edges and we use the phrase *network geometry* to indicate the distribution of edge weights [5]. The two candidate features that we consider are (i) the presence of cycles and (ii) the strength of connections in cycles. We will study these features through a rewiring process on an initial set of edges.

#### The presence of cycles

We begin by noting that cycles support temporally extended cascades. Given a single initial stimulus or spontaneous spike, a cascade can have a duration greater than the number of nodes in the graph if and only if there exists at least one cycle in the network. We demonstrate this simple intuition with an acyclic 3-node network and a cyclic 3-node network, where each edge in both networks has a weight of 0.5 (Figure 3a-b). In simulations of 10^4^ cascades, we find that the acyclic network produces a maximum cascade duration of 3 time steps, as expected. In contrast, using the same number of simulations on the cyclic network, we find the much greater maximum cascade duration of 13 time steps.

**Figure 3.**
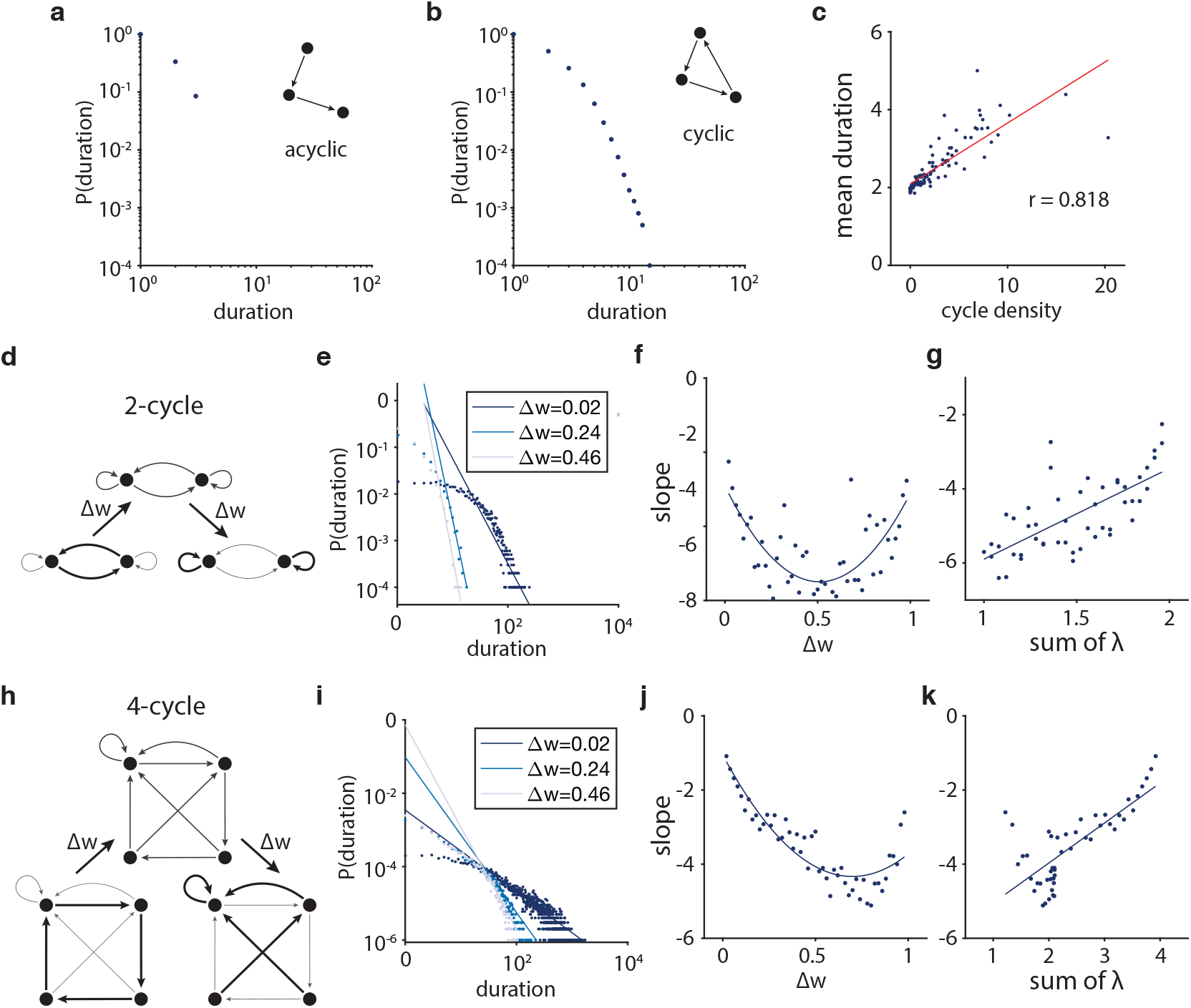
Cycles and strong connections support avalanching dynamics. **a**, The probability distribution of cascade duration in an acyclic network where all edges have a weight of 0.5. In 10^4^ trials, the maximum cascade duration is 3 time steps. **b**, The probability distribution of cascade duration in a cyclic network where all edges have a weight of 0.5. In 10^4^ trials, the maximum cascade duration is 13 time steps. **c**, Networks with higher cycle density have longer cascades. The cycle density is the number of simple cycles divided by the number of edges. A directed acyclic graph was randomly rewired to produce networks of varying cycle density. The Pearson’s correlation coefficient is *r =* 0.8180, *p =* 1.0835 x 10^-27^. **d**, A schematic of a 2-node network. The weights are redistributed from the 2-cycle to self-loops by Δ*ω*. **e**, Three probability distributions of cascade duration for Δ*ω* = 0.02, 0.24, and 0.46 in the 2-node cycle. The distributions have slopes of −2.2551, −5.6259, and −6.4050, respectively. **f**, The slope of distributions *(left)* decreases as Δ*ω* → 0.5 and *(right)* increases again as Δ*ω*→ 1. The quadratic fit of the slope of duration is *y =* 9.9509Δ*ω*^2^ - 10.1443Δ*ω* - 2.9108. **g**, Network geometries producing longer duration can be characterized by a larger sum of eigenvalues; note that these networks have the same dominant eigenvalues. The Pearson’s correlation coefficient is *r* = 0.6926, *p* = 3.5190 x 10^-8^. The y-axis is the slope of duration as in panel **f**. **h**, A schematic of a 4-node network. The weights are redistributed from the 4-cycle to randomly chosen edges that are not part of the original 4-cycle by Δ*ω*. **i**, Three probability distributions of cascade duration for Δ*ω*= 0.02, 0.24, and 0.46 in the 4-node cycle. The distributions have slopes of −1.0870, −2.6681, and −4.1164, respectively. **j**, The slopes of distributions follow the same shape as in panel **f**. The quadratic fit of the slope of duration is *y =* 6.7078Δ*ω*^2^ - 9.3683Δ*ω* - 1.0733. Note: The schematic (panel **h**) and simulations (panels **i**-**k**) use one random selection of edges that are not part of the original cycle. **k**, The 4-node cycle shows the same relationship between duration and sum of eigenvalues as in the 2-node cycle show in panel **g**. The Pearson’s correlation coefficient is *r* = 0.7611, *p* = 2.1791 x 10^-10^. The y-axis is the slope as in panel **j**.

Next, we show that the cascade duration scales monotically with a network’s cycle density, which we define as the number of simple cycles divided by the number of connected edges (Figure 3c). To study the effect of cycle density, we begin with a 10-node, directed acyclic graph and randomly rewire each edge with probability *p* to a different target node. The directed acyclic graph has the maximum number of edges; that is, the weight matrix is an upper triangular matrix without the diagonal entries. By sweeping over rewiring probabilities *p =* {0.0,0.1,0.2,…, 1.0}, we generate networks with different numbers of simple cycles, but the same number of edges and same edge weights of 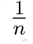. For each p, we simulate 10^4^ cascades with a maximum duration of 10^4^, and we measure the slope of the linear tail of the distribution on a log-log plot. In these simulations, we find that as a network is rewired to contain more cycles, the average cascade duration increases (Pearson’s correlation coefficient *r =* 0.8180, *p =* 1.0835 × 10^-27^). These examples illustrate the more general rule that networks containing cycles can support longer cascades and can extend the tail of the distribution of cascade duration.

#### The strength of connections in cycles

We now turn to a consideration of the distribution of edge weights. To maximize the specificity of our inferences and to generally build our intuition, we constrain ourselves initially to simple networks that only contain a small cycle (a 2-cycle) or that also contain one relatively larger cycle (a 4-cycle). We probe the role of weight distributions in the dynamics of the network by placing the strongest weights on edges comprising cycles and by placing the weakest weights on edges not comprising cycles. Specifically, in both the 2-node and 4-node cycle networks, we take the strong weights initially placed on the cycle and redistribute some of their weight by Δ*ω* to randomly chosen edges that are not part of the original cycle (Figure 3d,h). See the Supplementary Materials for results obtained for different random selection of edges. Upon these new networks, we simulate the stochastic model. We find that as the weight on the original cycle is continuously redistributed away from the initial cycle and throughout the network, we observe fewer and fewer cascades of long duration (Figure 3e,i).

Across empirical studies [6,7,16,23,41,49-51,56], the distributions of avalanche duration have been fit by power law functions, where the exponent is known as the life time. Typical values vary from −1.0 to −2.6. We seek to show how cycle density and edge weights in cycles together explain the topological and geometric differences in the networks underlying the various distributions of avalanche duration. The distribution of longer versus shorter cascades can be quantified by the slope of the power law distribution on a log-log plot. Cascades of longer duration would be expected to produce a shallower, or less negative, slope while cascades of shorter duration would be expected to produce a steeper, or less positive, slope. As we redistribute edge weight more uniformly in both the 2-node and 4-node networks, we find that the slopes of the distributions of cascade duration become increasingly negative, ranging from −2.26 to −6.41 for the 2-node network and from −1.09 to −5.11 for the 4-node network. Furthermore, as we redistribute away from the uniform geometry, continuously increasing the range of edge weights, we again observe more and more cascades of long duration (Figure 3f,j). These observations underscore the tight coupling between the range of edge weights, and the heavy-tailed nature of the distribution of cascade duration.

Lastly, we seek to determine whether the distribution of edge weights along cycles contributing to cascade duration is captured by the eigenvalue analysis presented in the previous section. Towards this goal, we employ the same perturbative numerical experiments on the 2-node and 4-node cycle networks. Specifically, we find that as the weights of the original cycle are redistributed, the duration of cascades, as measured by the slope of the distributions, tracks monotonically with the sum of eigenvalues of the network (Pearson’s correlation coefficients *r =* 0.6926, *p =* 3.5190 x 10^-8^ and *r* = 0.7611, *p =* 2.1791 x 10^-10^, respectively; Figure 3g,k). Because the dominant eigenvalues of the networks in these simulations are all equal to 1, the sums of eigenvalues provide a more descriptive estimation of activity propagation. Thus, this result suggests that edge weight constrains cascade duration by determining the strength of activity propagation.

### Node-specific dynamics

Even within a single network architecture, the range of cascade dynamics can vary depending on the nodes that are stimulated, either spontaneously or exogenously. Thus, we now consider the role of the stimulus pattern on cascade dynamics. We extend our eigenvalue analysis to estimate the role of a stimulus pattern on cascade dynamics by calculating the magnitude of the eigenprojection of the stimulus pattern. Because the average dynamics are explained by linear systems theory, we then use network control theory to more accurately predict how stimulation of individual nodes alters the dynamics of cascades.

#### The eigenprojection of the stimulus pattern

First, we tested whether the magnitude of the eigenprojection of the stimulus pattern could predict cascade dynamics. Given a stimulus ***y***(*0*), the eigendecomposition of the weight matrix *A* into *A = PDP^-1^* yields **c** = *P^-1^**y***(0) as the coefficients of the eigenmode excitation of *y*(0). The components of **c** determine how much the stimulus ***y***(0) projects onto the eigenvectors of *A* and describes the modes of average activity propagation through the network *A*. Then, as a predictor for mean duration, we can compute the 1-norm, or the sum of absolute values, of the eigenprojection of the stimulus pattern scaled by the corresponding eigenvalues ⋋,

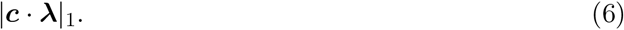

We numerically test the eigenprojection metric by simulating cascades on a 100-node, weighted random network. The mean duration of cascades generated from the stimulation of a single node was significantly positively correlated with the magnitude of the eigenprojection (Pearson’s correlation coefficient *r =* 0.4213, *p =* 1.27 x 10^-5^; Figure 4a). To determine the generalizability of these findings, we expanded our simulation set to include 30 random instantiations of networks with the same parameters. In this broader dataset, we found that the Pearson’s correlation coefficient was highly variable (median *r* = 0.1967; Figure 4d). Thus, we can weakly estimate the role of a stimulus pattern on cascade dynamics with eigenvalue analysis.

**Figure 4.**
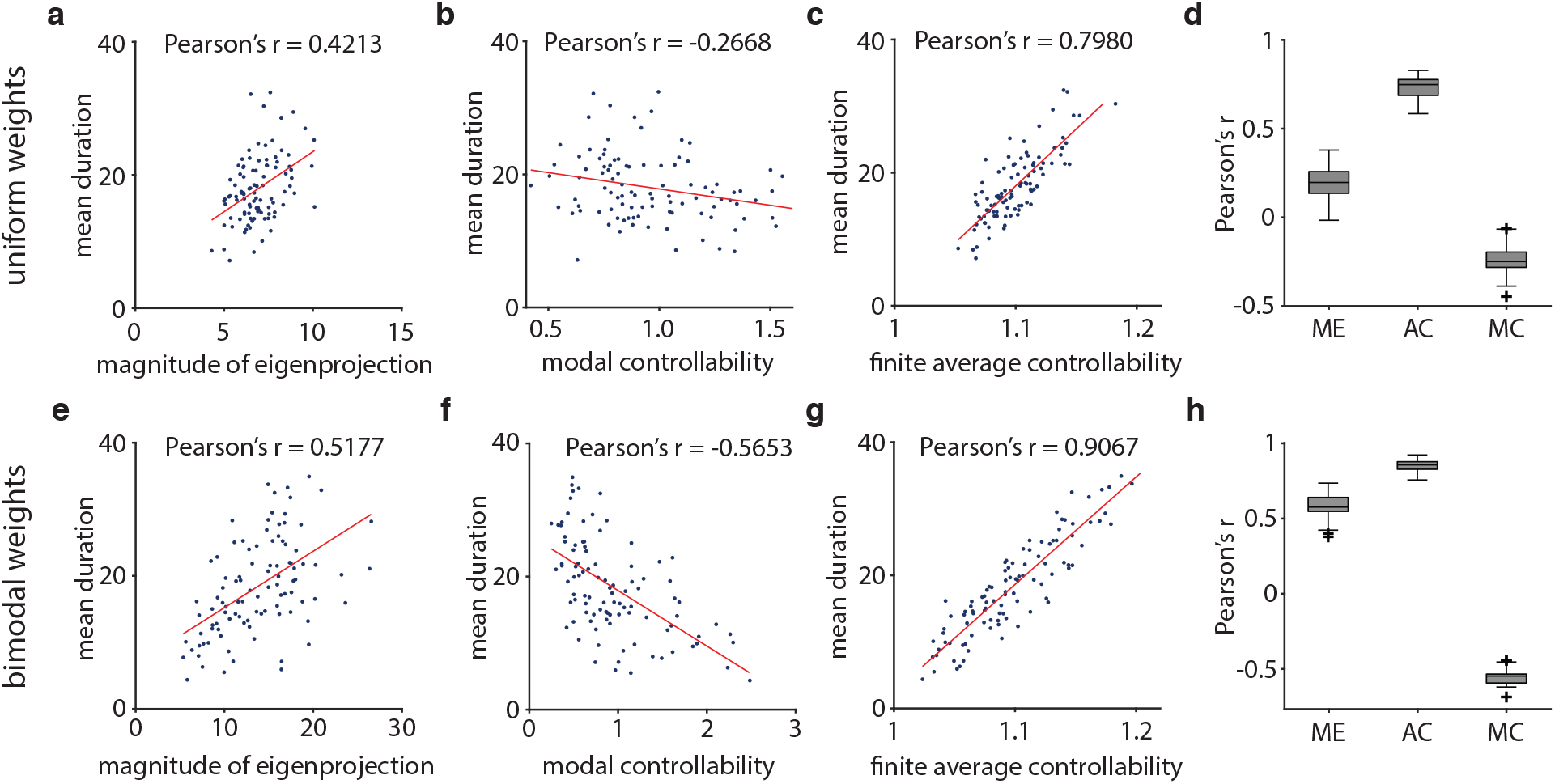
Network controllability is tightly linked with dynamics. **a-c**, Scatter plots between mean cascade duration and **a** the magnitude of the eigenprojection, **b** the modal controllability, and c the finite average controllability. The network used here has 100 nodes with weighted random graph topology and a fractional connectivity of around 0.2. Each point is the mean duration and metric for each node. **d**, Pearson’s correlation coefficients for 30 random instantiations of networks with the same parameters as the network used in panels **a-c**. **e-g**, Scatter plots between mean cascade duration and **e** the magnitude of the eigenprojection, **f** the modal controllability, and **g** the finite average controllability. The network used here has the same parameters as networks from panels **a-c**, but with 10% of connections normally distributed with a mean of 0.9 and 90% of connections with a mean of 0.1, all with a standard deviation of 0.1, before weight normalization. **h**, Pearson’s correlation coefficients for 30 random instantiations of networks with the same parameters as the network used in panels **e-g**.

#### Network control theory

To more accurately predict the role of a stimulus pattern on cascade dynamics, we adopt average and modal controllability metrics from linear control theory as we expect control of a network from a node to require network activity. In the same set of simulations reported above, we compared the mean cascade duration to the finite average controllability of each node, defined as

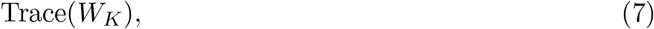

where 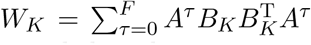 is the finite controllability Gramian (see Methods for details). We observed that the mean cascade duration and finite average controllability were significantly positively correlated (Pearson’s correlation coefficient *r =* 0.7980, *p =* 1.27 × 10^-5^; Figure 4c). In contrast, modal controllability was not strongly correlated with mean cascade duration (Pearson’s correlation coefficient *r =* −0.2668, *p =* 0.0073; Figure 4b). To determine the generalizability of these findings, we expanded our simulation set to include 30 random instantiations of networks with the same parameters. In this broader dataset, we observed consistent effects (median Pearson’s correlation coefficient *r =* 0.7483 and *r =* −0.2484 for finite average controllability and modal controllability, respectively; Figure 4d). In comparing the predictions from linear control theory with the predictions from eigendecomposition, we note that finite average controllability is consistently more strongly correlated with the mean cascade duration than the magnitude of the eigenprojection (Figure 4d).

Interestingly, networks with the same topological parameters as above, but with a bimodal distribution of weights show even stronger correlations between network control statistics and cascade dynamics (Figure 4f-h). Such a weight distribution reduces variance in the stochastic process, which intuitively can serve to strengthen the correlation. We observed that the mean cascade duration and finite average controllability were significantly positively correlated (Pearson’s correlation coefficient *r* = 0.9067, *p =* 1.6 x 10^-38^; Figure 4g). Modal controllability became strongly negatively correlated with mean cascade duration (Pearson’s correlation coefficient *r =* −0.5653, *p* = 8.9 x 10^-10^; Figure 4f). Again to determine the generalizability of these findings, we expanded our simulation set to include 30 random instantiations of networks with the same parameters. We observed consistent effects (median Pearson’s correlation coefficients between mean cascade duration and finite average controllability, modal controllability, and magnitude of eigenprojection were *r =* 0.8572, *r =* −0.5473, and *r =* 0.5765, respectively; Figure 4h). Again we note that finite average controllability is consistently more strongly correlated with the mean cascade duration than the magnitude of the eigenprojection (Figure 4h). These simulations suggest that the skewed weight distributions, as identified in the previous section as network motifs that support long cascades, may strengthen the relationship between network control and network dynamics. Collectively, the results illustrate that the stimulus patterns and the network must be tailored for each other to produce the desired neural dynamics. Our observations naturally lead to the question of how stimulation, either endogenous or exogenous, can be used for information processing.

### Cascade duration allows network discriminability and stimulus recovery

If certain network topologies and stimulus patterns can produce long-lasting cascades consistent with avalanche dynamics, what role can lasting cascades contribute to information processing? Intuitively, one cannot recover information about stimuli from cascades that have already terminated. For lasting cascades, network states can be discriminated and can also provide information about stimuli. Such delayed recovery of stimuli can allow the associative learning of stimuli across temporal delays [14,15,20]. The intuition that lasting cascades allow network discriminability can be formalized mathematically via Equation 3. Then, with simulations, we test the intuition that cascade dynamics support stimulus recoverability.

#### Network discriminability

To analytically show the relationship between cascade duration and discriminability, we first define network discriminability as the Euclidean distance between two states *d*(***y***_1_(*t*),***y***_2_(*t*)) in n-dimensional space. Recall that 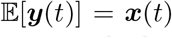 for stimulus *x*(0) from Equation 3. Then, given two stimuli, ***x***_1_(0) and ***x***_2_(0), we can calculate the expected network discriminability as the distance between the expected network states *d*(***x***_1_(*t*), ***x***_2_(*t*)) at time *t*. Given that the dominant eigenvalue ⋋_1_ < 1, then *x*(*t*) approaches the zero vector **0** as *t* approaches ∞. As described in previous sections, the decay in activity is constrained by the dominant eigenvalue of the network and by the finite average controllability of the individual node being stimulated. Thus, the rate at which both ***x***_1_(*t*) and ***x***_2_(*t*) decay to **0** determines the rate at which *d*(***x***_1_(*t*), ***x***_2_(*t*)) approaches *d*(**0, 0**) where discriminability between two network states is zero.

#### Stimulus recovery

To numerically show the relationship between cascade duration and stimulus recoverability, we first define stimulus recoverability as the mutual information *I*(*S;Y_t_*) between stimulus patterns *s* ∈ *S* and network states *y* ∈ *Y_t_* at time *t* (see Methods for details and Figure 5a-d for an intuitive schematic). Similar to discriminability, mutual information between the stimuli and network states decreases with shorter cascade duration because the Shannon entropy of the network states decreases. To probe this relation formally, we simulated cascades with 100-node networks from 4 different graph topologies with 30 instantiations of each graph type. Consistent with our intuition, we observe that mutual information is maintained longer when cascades last longer on average (Figure 5e-h). We then quantified the decay in mutual information by first performing linear regression on the mutual information as a function of time for the first 10 time steps. By calculating the Pearson’s correlation coefficient between the slope of linear regression and the mean cascade duration, we found that for all four graph topologies, mutual information decays faster when the propagation of activity also decays faster (Figure 5i-j). Collectively, these results demonstrate that stimulus recoverability is maintained longer when the cascades generated by stimulus patterns last longer.

To link information retention back to network structure, we assessed the relation between stimulus recoverability and the sum of eigenvalues of each network. Using the same 100-node networks from 4 different graph topologies with 30 instantiations of each graph type, we found a significant positive correlation between the average decay rate in mutual information and the sum of eigenvalues, implying that network structure supports the retention of information within the network (Pearson correlation coefficient *r* = 0.9188, *p* = 1.8 × 10^-49^; Figure 5k). Moreover, while all networks had similar parameters, each graph type generated distinct ranges of decay rates and sums of eigenvalues, suggesting that certain graph types may be better suited for information retention than others. In particular, we observed lower decay rates in mutual information and lower sum of eigenvalues in the weighted random and modular graphs, than in the random geometric and Watts-Strogatz graphs. Collectively, these findings demonstrate the interplay among network architecture, network dynamics, and information processing.

## Discussion

Neural systems display strikingly rich dynamics that harbor the marks of a complex underlying network architecture among units, from the small scale of individual neurons to the large scale of columns and areas [46, 76]. Avalanches are a quintessential example of such dynamics, and are thought to allow for a diverse range of computations [6, 24, 36, 57]. Yet, precisely how a neuronal network’s topology supports stochastic dynamics and the computations that can arise therefrom remains unclear. Here, we seek to provide clarity using both precise analysis of mathematical formulations and statistically rigorous assessments of numerical experiments. We consider a generalized spiking model and demonstrate that the time-averaged activity of this model can be treated as a linear dynamical system. From this observation, we derive intuitions for how network topology constrains cascade duration. In subsequent numerical experiments, we use eigendecomposition and statistical approaches from network control theory appropriate for linear dynamical systems to describe how network structure and the stimulus pattern together determine the manner in which a stimulus propagates through the network. We identify strongly connected cycles as prevalent network motifs that promote long cascade duration in neuronal networks. Finally, we use mutual information to demonstrate that long-lasting cascades can serve as a mechanism to allow for temporally delayed recovery of desired patterns of stimulation. Broadly, our work blends dynamical systems theory, network control theory, information theory, and computational neuroscience to address the wide gap in the field’s current understanding of the relations between architecture, dynamics, and computation.

**Figure 5.**
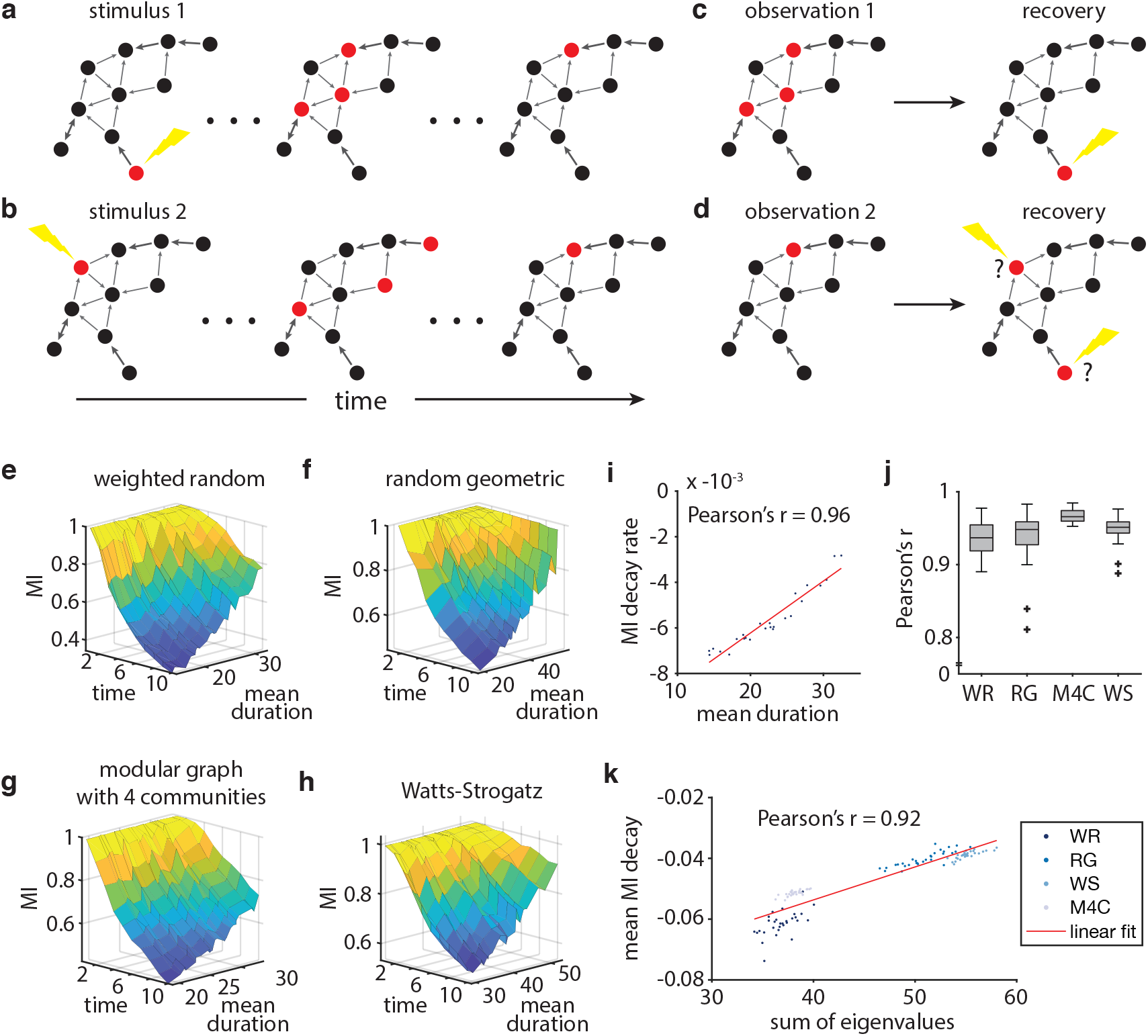
A stimulus can be well-recovered when it generates long-lasting cascades. **a-b**, A schematic showing two cascades triggered by different stimuli. **c**, Recovery of the stimulus using an observation of a network state during a cascade. **d**, Failed recovery of the stimulus using an observation of a network state during a cascade. **e-h**, Decay in mutual information (MI) over time. When activity from a stimulus pattern lasts longer, mutual information also persists for longer. Panels **e-h** show results from four graph types: a weighted random graph, a random geometric graph, a modular graph with 4 communities, and a Watts-Strogatz graph. We used a fractional connectivity of 0.05 to show a wide range of decay rates in mutual information (see Supplemental Materials for simulations with other fractional connectivities). **i**, The linear decay rate of mutual information over the first 10 time steps plotted against the mean cascade duration in the example weighted random graph from panel **e**. **j**, A boxplot of the Pearson correlation coefficients between the linear slope of decay in mutual information over time and the mean cascade duration. The boxplot shows data from 30 instantiations of each graph type, each network containing 100 nodes and characterized by a fractional connectivity of around 0.05. **k**, The mean decay rate in mutual information for a network is correlated with the sum of eigenvalues of the network (Pearson’s correlation coefficient *r* = 0.9188, *p* = 1.8 × 10^−49^).

### Linear form of stochastic network dynamics

Because of the inherently stochastic nature of neuronal avalanches, many previous studies have simply inferred properties about the underlying network through statistical methods [6, 40]. An important innovation in this study was the demonstration that the time-averaged activity of the stochastic system has an equivalent form as a linear dynamical system. Such linear estimation of the dynamics makes available powerful computational tools in matrix and linear systems theory, and allowed us to capitalize on recent advances in network control [39, 47]. Network control theory is a formal approach to modeling, predicting, and tuning the response of a networked system to exogenous input, and has been recently applied to neural systems at both the cellular [73, 78, 80] and regional [13, 22, 31, 67] scales (for a recent review, see [66]). In these previous efforts, linear dynamics have been assumed, whereas here such dynamics have been proven, to be relevant for the neural system under study. Extensions of linear systems analysis, such as observability [12] and optimal control [10, 21, 68], follow immediately from this work and could provide added insights into other dynamical and computational properties of neural networks. Finally, it would be of interest to directly probe the effects of stimulation patterns defined by network controllability statistics on information transmission in vitro or behaviors in vivo, following work in a similar vein in large-scale human neuroimaging [34, 43, 45, 65].

### Topological constraints on dynamics and computation

Proving formally that network topology affects dynamics and computation is important, but can be further complemented by providing intuitions regarding the specific features of a network topology that are most relevant. The identification of functionally relevant features of networked systems has a long history in molecular biology [2], with notable efforts identifying structural motifs in transcription regulation networks [55], protein-protein interaction networks [81], and cellular circuits [26], which are thought to arise spontaneously under evolutionary pressures [33]. Significantly extending prior statistical efforts in large-scale connectomes [64], here we demonstrate that specific structural motifs in the form of strongly connected cycles are topological features that support long cascade dynamics. These structural motifs form elementary units or building blocks of the network that can be combined to create connectivity architectures that produce certain dynamical behaviors. Other theoretical studies have also found strongly and bi-directionally connected neurons as motifs that produce long-lasting memory [11], potentially as a mechanism for attractor dynamics [30]. Importantly, empirical studies have shown that the network motifs identified here are observed in both cortical microcircuits [37, 38, 42, 48, 63, 75] and macrocircuits [60]. Future work is needed to better understand the rules by which neurons connect to one another, and to determine whether those rules serve to increase the memory capacity of cortical networks. It would also be interesting in the future to determine whether higher-order structural motifs, such as those accessible to tools from algebraic topology [19, 61], might also play a role in the relationships between topology, dynamics, and computation [53, 60].

### Information theory as a performance measure

To measure information retention, we use mutual information between stimulus patterns and network states. Mutual information, originally developed to study communication channels [54], has proven to be a powerful tool for the study of information transmission in avalanching neural networks [6, 58]. While previous studies of neuronal avalanches use power law statistics that suggest criticality as the theoretical link between dynamics and information processing [6, 9, 24, 36, 56, 57, 59], we take a more mechanistic approach embedded in dynamical systems theory to study the relationships between network structure, dynamics, and mutual information. In light of recent evidence for the subcriticality of cortical networks and the difficulty of establishing criticality from power laws [52, 71, 72], the direct approach that we take here may prove useful in future studies. Nevertheless, we also acknowledge that our approach has some limitations. Despite its utility in studying information channels, mutual information is unlikely to be the only useful performance measure for a neural system, given the numerous purported computations of cortical networks [57, 70]. Indeed, the explanation posited here for the prevalence of strongly connected neurons does not account for the information faculties of the rest of the neural system. Such considerations compel further investigation into how network structure supports other types of information processing accessible to other information theoretic measures.

### Methodological considerations

A few remarks are warranted on the topic of linear dynamics in neural systems. Linear dynamics accurately predicts stochastic, cascade dynamics, and its rich mathematical properties have been used to study neural dynamics in many organisms across a wide range of temporal and spatial scales [22, 35, 39, 80]. At the neuronal level, however, neural dynamics are non-linear [28]. Efforts analytically demonstrating properties about non-linear systems are more limited [44], and thus, further study is required to more thoroughly demonstrate the relationships shown here in a non-linear system.

### Future directions

In closing, we note that the natural direction in which to take this work will be to consider other types of information processing and to identify network structures that produce dynamics that support such computations. Moreover, it would be apt to apply this framework to cortical networks from functional, structural, and effective connectivities and to measure memory performance in terms of the network topology and dynamics. It would be interesting to measure differences in memory performance across brain regions, and to test for relationships between topological features and performance. Third and finally, studying well-known network learning rules—such as Hebbian plasticity [27] and spike-timing dependent plasticity [62]—in a dynamical systems and information theoretic framework may shed further light on the functional purpose of these rules.

## Acknowledgments

H.J., J.Z.K., and D.S.B. acknowledge support from the John D. and Catherine T. MacArthur Foundation, the Alfred P. Sloan Foundation, the ISI Foundation, the Paul Allen Foundation, the Army Research Laboratory (W911NF-10-2-0022), the Army Research Office (Bassett-W911NF-14-1-0679, Grafton-W911NF-16-1-0474, DCIST-W911NF-17-2-0181), the Office of Naval Research, the National Institute of Mental Health (2-R01-DC-009209-11, R01-MH112847, R01-MH107235, R21-M MH-106799), the National Institute of Child Health and Human Development (1R01-HD086888-01), National Institute of Neurological Disorders and Stroke (R01-NS099348), and the National Science Foundation (BCS-1441502, BCS-1430087, NSF PHY-1554488 and BCS-1631550). J.Z.K acknowledges support from the NIH T32-EB020087, PD: Felix W. Wehrli, and the National Science Foundation Graduate Research Fellowship No. DGE-1321851. We thank Erin Teich, Lia Papadopoulos, and Zhixin Lu for comments and suggestions on the paper. The content is solely the responsibility of the authors and does not necessarily represent the official views of any of the funding agencies.

## Methods

### Network generation

We study a recurrent, excitatory network of *n* neurons connected by connectivity matrix *A*. A network is formally represented by directed graph 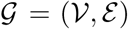, where 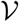 = {1,…, *n*}, and *ε* ⊆ 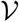 x 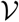 are the sets of network vertices and directed edges. Let *A* = [*a_ij_*] be the weighted and directed adjacency matrix of 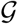, where *a_ij_* is the connection from neuron *j* to neuron *i* and 0 ≤ *a_ij_* ≤ 1. At each time point *t* ∊ ℤ_≥0_, we associate each node *i* with a discrete non-negative random variable *y_i_(t)*.

We use four different commonly studied graph models from network science in our analyses [79]. The first graph model is the *Weighted Random Graph* model (WRG), which is a weighted version of the canonical Erdos-Renyi model. The weight of an edge is distributed as a geometric distribution with probability of success *p*. Second, we use a *Random Geometric* model (RG) that is embedded in a unit cube, where the edge weights are equal to the inverse of the Euclidean distance between two nodes. We kept only a fraction of the shortest edges in order to achieve a desired edge density *p*. Third, we use a *Modular Graph with 4 Communities* model (MD4). Pairs of nodes within communities have an edge density of 0.8, and nodes across communities are connected to achieve a desired edge density of *p*. The edges of nodes in the same community and across communities are weighted according to a geometric distribution with probability of success *p* and 1 – *p*, respectively. Fourth, we use a *Watts-Strogatz* model (WS). The model builds a ring lattice and then uniformly rewires the network, creating a small-world architecture with a random probability of *r* = 0.1. After a graph is generated, we normalize the weights so that the sum of each row is equal to 1.

To calculate the cycle density of a graph, we compute the number of simple cycles divided by the number of connected edges. A simple cycle is defined as the set of edges in a closed walk with no repetitions of vertices and edges, other than the starting and ending vertex. The number of simple cycles was calculated using the NetworkX software package (version 2.1) on python v2.7.10.

### Stochastic McCulloch-Pitts model

We model cascades as spikes propagating through a recurrent network. To model neuronal cascades, we next stipulate a stochastic version of the McCulloch-Pitts neuron. In the McCulloch-Pitts model, a neuron receives inputs scaled by the weights of the edges and sums the scaled inputs to produce an output via an activation function. Here, the activation function is a Bernoulli process, where probability *p* is the sum of the scaled inputs. The model is similar to the branching model that has been applied to networks to study neuronal avalanches [6, 24, 25, 52]. The network starts at some non-random initial state ***y***(0), which can also be interpreted as a stimulus received at *t* = 0. Activity in the form of binary spikes {0,1} propagates through the network as ***y**(t)*. For neuron *i*, *y_i_(t)* is defined as

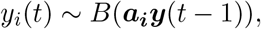

where *B(p)* is a Bernoulli process with probability *p*, *a_ij_* is the weight from neuron *j* to neuron *i*, and 0 ≤ *a_ij_* ≤ 1. For computational tractability, we set a maximum time step *K* for the simulations. The simulated spike counts ***y**(t)* are stored as a *n*-by-*K* matrix. All simulations and calculations were run on MATLAB (version 2018a) provided by The MathWorks, Inc.

We prove by induction that linear dynamics estimates average stochastic behavior, i.e., 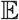[*y_j_(t)*] = *x_j_(t)*. At *t* = 0, both *y_j_*(0) and *x_j_*(0) are set as the stimulus pattern, and so, 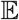[*y_j_*(0)] = *x_j_*(0). Now, assume 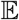[*y_j_*(*t* – 1)] = *x_j_*(*t* – 1), and see that 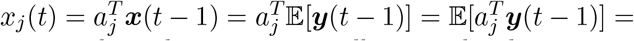 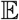[*y_j_(t)*] and thus, 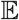[*y_j_(t)*] = *x_j_(t)*. To demonstrate this relation numerically, we take the average cascades that begin with the same initial state by taking the mean of 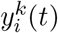 for all cascades *k* at each time step *t*. All cascades start with the same initial condition *y(0)*.

**Eigenvalue analysis.** In our analysis of networks, we decompose the weight matrix *A* into eigenvalues and eigenvectors. Such an eigendecomposition is formalized as

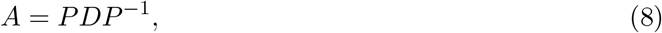

where *P* is a matrix of eigenvectors as columns and D is a diagonal matrix of corresponding eigenvalues. According to the Perron-Frobenius theorem, the real, square, and non-negative adjacency matrices used here all have a largest real eigenvalue, and the corresponding eigenvector can be chosen to have strictly non-negative components.

### Stimulus pattern generation

We investigate the propagation of activity through a network initiated by stimulus patterns. The stimulus pattern is set as the initial state *y*(0) or *x*(0) of a network and then propagated forward in time according to either a branching process or linear dynamics, respectively. In our study, we consider two ways to generate stimulus patterns. In the analysis of cascade duration and controllability, we stimulate individual nodes by creating a set of vectors in which the *i^th^* element of the *i^th^* vector is set at 1 and all other elements are set at 0. In the mutual information analysis, we create a set of column vectors such that their finite average controllability values evenly span the range of controllability values (see later section of this Methods for definition of finite average controllability). In each of the 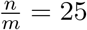 vectors, we choose *m* = 4 nodes from *n* = 100 total nodes to stimulate such that each node that we select is increasing in its finite average controllability value. Because finite average controllability is highly correlated with cascade duration, such input vectors will evenly span the possible duration of cascades.

### Predicting cascade dynamics

We can predict the exact fraction of cascades alive at time *t* by computing a state transition matrix from any state *k* to any state *l*. For *k, l* ∊ {1,…, *n*}, the state transition matrix *T* ∊ ℝ^*l × k*^ can be constructed by

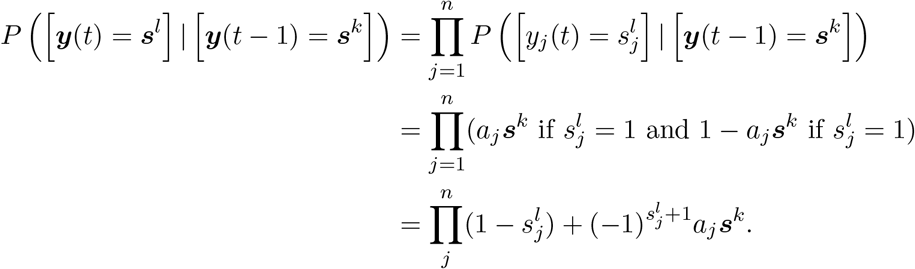

Then, at *t* for all *l*, the probability of the network being in any state is given by *P(y(t) = s^l^)* = 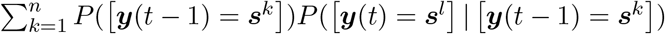.

### Network control theory and associated controllability statistics

Network control theory is a formulation of control theory for networks of interacting components. This formulation typically consists of a set of *n* component nodes 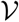 = {1,…, *n*}, where the vector ***x**(t)* ∊ ℝ^*n*^ represents the state of node activities at time *t* ≥ 0. These nodes are connected by a set of edges *ε* ⊆ 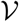 × 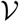, where the adjacency matrix *A* ∊ ℝ^*n × n*^ has elements *a_ij_* as the strength of the connection from node *j* to node *i*. Here, *control* typically refers to a set of *k* inputs *u(t)* ∊ ℝ^k^ at time *t* ≥ 0 that drive the evolution of system states according to *B* ∊ ℝ^*n × k*^. In linear control theory, the system states evolve as

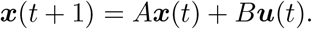

Motivated by a desire to understand how network architecture affects its control properties, recent work iterates network-based metrics for control of such linear systems [47]. Particularly germane to our discussion of avalanche duration is *average controllability* [22, 32], defined as the H2 norm of the system’s infinite impulse response given by

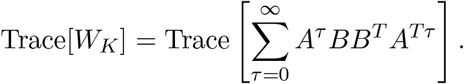

Here, we set *B* as a binary column vector where vector elements corresponding to the nodes of interest are set to 1 and the remaining vector elements are set to 0; this formulation represents an impulse of magnitude 1 to the nodes of interest. The finite average controllability is similarly defined by taking the sum to some finite positive integer *F* instead of infinity, and represents the norm of the system’s impulse response over *F* time steps. Because avalanches are expected to last for a finite number of time steps, we use *F* = 10 in the main text, and in the supplement we show that larger values of F produce similar results.

Another network-based control metric we use here is *modal controllability* [22, 47]. While modal controllability was originally formulated for symmetric matrices, here we extend the definition to include asymmetric matrices. To do this, we take the absolute value of both the eigenvalues and the eigenvector components, which can be complex numbers in an asymmetric matrix. Thus, we define the version of modal controllability of node *i* for asymmetric matrices as

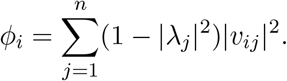

### Mutual information calculation to probe stimulus recovery

To measure the capacity of a network to transfer information during an avalanche, we calculated the mutual information *I(X; Y)*, which quantifies the amount of information, in bits, that one random variable *X* reveals about another random variable *Y*. Here, the two random variables of interest are the initial stimulus patterns ***y***(0) = ***s*** ∊ *S* and the measured network states ***y**(t)* ∊ *Y_t_* at each time step *t*. With mutual information, we measure the amount of information that the network states *Y_t_* reveal about which stimulus pattern ***y***(0) ∊ ***s**_i_* or *S′* was presented. To obtain a reasonable set of stimulus patterns, we generated the set *S′* = {***s**_j_*|*j* ≠ *i*} of |*S*′| = *n* – 1 unique stimulus patterns. Both *P*(***y***(0) = ***s**_i_*) and *P*(***y***(0) ∊ *S*′) are equally probable at 0.5 such that *H*(***y***(0)) ≈ 1 bit for all patterns *i*. We generated the stimulus patterns such that they evenly spanned the range of average controllability values (see earlier Methods section on “Stimulus pattern generation”). All mutual information calculations were run using the MIToolbox (v3.0.1) for MATLAB (https://github.com/Craigacp/MIToolbox).

In the analysis of the relationship between the average cascade duration and the mutual information, we quantify the decay in mutual information over time. We also calculate the correlation between the decay rate of the mutual information and the predicted mean cascade duration. For this latter calculation, first we perform a linear regression of the decay in mutual information with respect to time. Then, we calculate the Pearson correlation coefficient between the slope of the linear regression and the mean cascade duration.

## References

1. R. Adolphs. Cognitive neuroscience of human social behaviour. Nature Reviews Neuroscience, 4:165 EP –, 03 2003.

2. U. Alon. Network motifs: theory and experimental approaches. Nat Rev Genet, 8(6):450–461, 2007.

3. D. J. Amit and N. Brunel. Model of global spontaneous activity and local structured activity during delay periods in the cerebral cortex. Cereb Cortex, 7(3):237–252, Apr-May 1997.

4. P. Bak, C. Tang, and K. Wiesenfeld. Self-organized criticality: An explanation of the 1/f noise. Physical Review Letters, 59(4):381–384, July 1987.

5. D.S. Bassett, M. A. Porter, N. F. Wymbs, S. T. Grafton, J. M. Carlson, and P. J. Mucha. Robust detection of dynamic community structure in networks. Chaos, 23(1):013142, Mar 2013.

6. J. M. Beggs and D. Plenz. Neuronal avalanches in neocortical circuits. Journal of Neuroscience, 23(35):11167–11177, 2003.

7. T. Bellay, A. Klaus, S. Seshadri, and D. Plenz. Irregular spiking of pyramidal neurons organizes as scale-invariant neuronal avalanches in the awake state. eLife, 4:e07224, jul 2015.

8. R. Ben-Yishai, R. L. Bar-Or, and H. Sompolinsky. Theory of orientation tuning in visual cortex. Proceedings of the National Academy of Sciences, 92(9):3844–3848, 1995.

9. N. Bertschinge. and T. Natschläger. Real-time computation at the edge of chaos in recurrent neural networks. Neural Computation, 16(7):1413–1436, 2004.

10. R. F. Betzel, S. Gu, J. D. Medaglia, F. Pasqualetti, and D. S. Bassett. Optimally controlling the human connectome: the role of network topology. Sci Rep, 6:30770, 2016.

11. N. Brunel. Is cortical connectivity optimized for storing information? Nature Neuroscience, 19:749 EP –, 04 2016.

12. C.-T. Chen,. Linear System Theory and Design. Oxford University Press, Inc., New York, NY, USA, 3rd edition, 1998.

13. E. J. Cornblath, E. Tang, G. L. Baum, T. M. Moore, A. Adebimpe, D. R. Roalf, R. C. Gur, R. E. Gur, F. Pasqualetti, T. D. Satterthwaite, and D. S. Bassett. Sex differences in network controllability as a predictor of executive function in youth. Neuroimage, 188:122–134, 2018.

14. D. Durstewitz, J. K. Seamans, and T. J. Sejnowski. Neurocomputational models of working memory. Nature Neuroscience, 3:1184 EP –, 11 2000.

15. J. Eriksson, E. K. Vogel, A. Lansner, F. Bergström, and L. Nyberg. Neurocognitive architecture of working memory. Neuron, 88(1):33–46, 2015.

16. N. Friedman, S. Ito, B. A. W. Brinkman, M. Shimono, R. E. L. DeVille, K. A. Dahmen, J. M. Beggs, and T. C. Butler. Universal critical dynamics in high resolution neuronal avalanche data. Phys. Rev. Lett., 108:208102, May 2012.

17. D. Garlaschelli. The weighted random graph model. New Journal of Physics, 11(7):073005, 2009.

18. E. D. Gireesh and D. Plenz. Neuronal avalanches organize as nested theta- and beta/gamma-oscillations during development of cortical layer 2/3. Proceedings of the National Academy of Sciences, 105(21):7576–7581, 2008.

19. C. Giusti, R. Ghrist, and D. S. Bassett. Two’s company, three (or more) is a simplex: Algebraic-topological tools for understanding higher-order structure in neural data. J Comput Neurosci, 41(1):1–14, 2016.

20. P. Goldman-Rakic. Cellular basis of working memory. Neuron, 14(3):477 – 485, 1995.

21. S. Gu, R. F. Betzel, M. G. Mattar, M. Cieslak, P. R. Delio, S. T. Grafton, F. Pasqualetti, and D. S. Bassett. Optimal trajectories of brain state transitions. Neuroimage, 148:305–317, 2017.

22. S. Gu, F. Pasqualetti, M. Cieslak, Q. K. Telesford, A. B. Yu, A. E. Kahn, J. D. Medaglia, J. M. Vettel, M. B. Miller, S. T. Grafton, and D. S. Bassett. Controllability of structural brain networks. Nature Communications, 6:8414 EP –, 10 2015.

23. G. Hahn, T. Petermann, M. N. Havenith, S. Yu, W. Singer, D. Plenz, and D. Nikolić. Neuronal avalanches in spontaneous activity in vivo. Journal of Neurophysiology, 104(6):3312–3322, 2010. PMID: 20631221.

24. C. Haldema. and J. M. Beggs. Critical branching captures activity in living neural networks and maximizes the number of metastable states. Phys. Rev. Lett., 94:058101, Feb 2005.

25. T. E. Harris. The theory of branching processes. The Rand Corporation, May 1964.

26. Y. Hart, Y. E. Antebi, A. E. Mayo, N. Friedman, and U. Alon. Design principles of cell circuits with paradoxical components. Proc Natl Acad Sci U S A, 109(21):8346–8351, 2012.

27. D. Hebb. The Organization of Behavior: a Neuropsychological Theory. Oxford, England: Wiley, 1949.

28. A. L. Hodgkin and A. F. Huxley. A quantitative description of membrane current and its application to conduction and excitation in nerve. The Journal of physiology, 117(4):500–544, 08 1952.

29. C. J. Honey, R. Kötter, M. Breakspear, and O. Sporns. Network structure of cerebral cortex shapes functional connectivity on multiple time scales. Proceedings of the National Academy of Sciences, 104(24):10240–10245, 2007.

30. J. J. Hopfield. Neural networks and physical systems with emergent collective computational abilities. Proceedings of the National Academy of Sciences, 79(8):2554–2558, 1982.

31. J. Jeganathan, A. Perry, D. S. Bassett, G. Roberts, P. B. Mitchell, and M. Breakspear. Fronto-limbic dysconnectivity leads to impaired brain network controllability in young people with bipolar disorder and those at high genetic risk. Neuroimage Clin, 19:71–81, 2018.

32. T. Kailath. Linear Systems. Prentice-Hall, 1980.

33. N. Kashta. and U. Alon. Spontaneous evolution of modularity and network motifs. Proc Natl Acad Sci U S A, 102(39):13773–13778, 2005.

34. A. N. Khambhati, A. E. Kahn, J. Costantini, Y. Ezzyat, E. A. Solomon, R. E. Gross, B. C. Jobst, S. A. Sheth, K. A. Zaghloul, G. Worrell, S. Seger, B. C. Lega, S. Weiss, M. R. Sperling, R. Gorniak, S. R. Das, J. M. Stein, D. S. Rizzuto, M. J. Kahana, T. H. Lucas, K. A. Davis, J. I. Tracy, and D. S. Bassett. Predictive control of electrophysiological network architecture using direct, single-node neurostimulation in humans. bioRxiv, 292748, 2018.

35. J. Z. Kim, J. M. Soffer, A. E. Kahn, J. M. Vettel, F. Pasqualetti, and D. S. Bassett. Role of graph architecture in controlling dynamical networks with applications to neural systems. Nature Physics, 14:91–98, 09 2018.

36. O. Kinouch. and M. Copelli. Optimal dynamical range of excitable networks at criticality. Nature Physics, 2:348 EP –, 04 2006.

37. H. Ko, S. B. Hofer, B. Pichler, K. A. Buchanan, P. J. Sjöström, and T. D. Mrsic-Flogel. Functional specificity of local synaptic connections in neocortical networks. Nature, 473:87 EP –, 04 2011.

38. S. Lefort, C. Tomm, J.-C. F. Sarria, and C. C. Petersen. The excitatory neuronal network of the c2 barrel column in mouse primary somatosensory cortex. Neuron, 61(2):301 – 316, 2009.

39. Y.-Y. Liu, J.-J. Slotine, and A.-L. Barabási. Controllability of complex networks. Nature, 473:167 EP –, 05 2011.

40. F. Lombardi, H. J. Herrmann, C. Perrone-Capano, D. Plenz, and L. de Arcangelis. Balance between excitation and inhibition controls the temporal organization of neuronal avalanches. Phys. Rev. Lett., 108:228703, May 2012.

41. F. Lombardi, H. J. Herrmann, D. Plenz, and L. D. Arcangelis. On the temporal organization of neuronal avalanches. Frontiers in Systems Neuroscience, 8:204, 2014.

42. H. Markram. A network of tufted layer 5 pyramidal neurons. Cerebral Cortex, 7(6):523–533, 1997.

43. J. D. Medaglia, D. Y. Harvey, N. White, A. Kelkar, J. Zimmerman, D. S. Bassett, and R. H. Hamilton. Network controllability in the inferior frontal gyrus relates to controlled language variability and susceptibility to TMS. J Neurosci, 38(28):6399–6410, 2018.

44. A. E. Motter. Networkcontrology. Chaos (Woodbury, N.Y.), 25(9):097621; 097621–097621, 09 2015.

45. S. F. Muldoon, F. Pasqualetti, S. Gu, M. Cieslak, S. T. Grafton, and D. S. Vettel, J M ad Bassett. Stimulation-based control of dynamic brain networks. PLoS Comput Biol, 12(9):e1005076, 2016.

46. S. Nigam, M. Shimono, S. Ito, F.-C. Yeh, N. Timme, M. Myroshnychenko, C. C. Lapish, Z. Tosi, P. Hottowy, W. C. Smith, S. C. Masmanidis, A. M. Litke, O. Sporns, and J. M. Beggs. Rich-club organization in effective connectivity among cortical neurons. Journal of Neuroscience, 36(3):670–684, 2016.

47. F. Pasqualetti, S. Zampieri, and F. Bullo. Controllability metrics, limitations and algorithms for complex networks. IEEE Transactions on Control of Network Systems, 1(1):40–52, 2014.

48. R. Perin, T. K. Berger, and H. Markram. A synaptic organizing principle for cortical neuronal groups. Proceedings of the National Academy of Sciences, 108(13):5419–5424, 2011.

49. T. Petermann, T. C. Thiagarajan, M. A. Lebedev, M. A. L. Nicolelis, D. R. Chialvo, and D. Plenz. Spontaneous cortical activity in awake monkeys composed of neuronal avalanches. Proceedings of the National Academy of Sciences, 106(37):15921–15926, 2009.

50. S.-S. Poil, R. Hardstone, H. D. Mansvelder, and K. Linkenkaer-Hansen. Critical-state dynamics of avalanches and oscillations jointly emerge from balanced excitation/inhibition in neuronal networks. Journal of Neuroscience, 32(29):9817–9823, 2012.

51. A. Ponce-Alvarez, A. Jouary, M. Privat, G. Deco, and G. Sumbre. Whole-brain neuronal activity displays crackling noise dynamics. Neuron, 11 2018.

52. V. Priesemann, M. Wibral, M. Valderrama, R. Pröpper, M. L. Van Quyen, T. Geisel, J. Triesch, D. Nikolić, and M. H. J. Munk. Spike avalanches in vivo suggest a driven, slightly subcritical brain state. Frontiers in Systems Neuroscience, 8:108, 2014.

53. M. W. Reimann, M. Nolte, M. Scolamiero, K. Turner, R. Perin, G. Chindemi, P. Dlotko, R. Levi, K. Hess, and H. Markram. Cliques of neurons bound into cavities provide a missing link between structure and function. Front Comput Neurosci, 11:48, 2017.

54. C. E. Shannon. A mathematical theory of communication. Bell System Technical Journal, 27(3):379–423, 1948.

55. S. S. Shen-Orr, R. Milo, S. Mangan, and U. Alon. Network motifs in the transcriptional regulation network of Escherichia coli. Nat Genet, 31(1):64–68, 2002.

56. W. L. Shew, W. P. Clawson, J. Pobst, Y. Karimipanah, N. C. Wright, and R. Wessel. Adaptation to sensory input tunes visual cortex to criticality. Nature Physics, 11:659 EP –, 06 2015.

57. W. L. Shew, H. Yang, T. Petermann, R. Roy, and D. Plenz. Neuronal avalanches imply maximum dynamic range in cortical networks at criticality. Journal of Neuroscience, 29(49):15595–15600, 2009.

58. W. L. Shew, H. Yang, S. Yu, R. Roy, and D. Plenz. Information capacity and transmission are maximized in balanced cortical networks with neuronal avalanches. Journal of Neuroscience, 31(1):55–63, 2011.

59. O. Shriki, J. Alstott, F. Carver, T. Holroyd, R. N. Henson, M. L. Smith, R. Coppola, E. Bullmore, and D. Plenz. Neuronal avalanches in the resting meg of the human brain. Journal of Neuroscience, 33(16):7079–7090, 2013.

60. A. E. Sizemore, C. Giusti, A. Kahn, J. M. Vettel, R. F. Betzel, and D. S. Bassett. Cliques and cavities in the human connectome. J Comput Neurosci, 44(1):115–145, 2018.

61. A. E. Sizemore, J. E. Phillips-Cremins, R. Ghrist, and D. S. Bassett. The importance of the whole: Topological data analysis for the network neuroscientist. Network Neuroscience, Epub Ahead of Print, 2018.

62. S. Song, K. D. Miller, and L. F. Abbott. Competitive hebbian learning through spike-timing-dependent synaptic plasticity. Nature Neuroscience, 3:919 EP –, 09 2000.

63. S. Song, P. J. Sjöström, M. Reigl, S. Nelson, and D. B. Chklovskii. Highly nonrandom features of synaptic connectivity in local cortical circuits. PLOS Biology, 3(3), 03 2005.

64. O. Sporn. and R. Kotter. Motifs in brain networks. PLoS Biol, 2(11):e369, 2004.

65. J. Stiso, A. N. Khambhati, T. Menara, A. E. Kahn, J. M. Stein, S. R. Das, R. Gorniak, J. I. Tracy, B. Litt, K. A. Davis, T. Pasqualetti, T. H. Lucas, and D. S. Bassett. White matter network architecture guides direct electrical stimulation through optimal state transitions. arXiv, 1805:01260, 2018.

66. E. Tan. and D. S. Bassett. Control of dynamics in brain networks. Rev. Mod. Phys., 90:031003, 2018.

67. E. Tang, C. Giusti, G. L. Baum, S. Gu, E. Pollock, A. E. Kahn, D. R. Roalf, T. M. Moore, K. Ruparel, R. C. Gur, R. E. Gur, T. D. Satterthwaite, and D. S. Bassett. Developmental increases in white matter network controllability support a growing diversity of brain dynamics. Nat Commun, 8(1):1252, 2017.

68. P. N. Taylor, J. Thomas, N. Sinha, J. Dauwels, M. Kaiser, T. Thesen, and J. Ruths. Optimal control based seizure abatement using patient derived connectivity. Front Neurosci, 9:202, 2015.

69. C. Tetzlaff, S. Okujeni, U. Egert, F. Wörgötter, and M. Butz. Self-organized criticality in developing neuronal networks. PLOS Computational Biology, 6(12):1–18, 12 2010.

70. N. M. Timme, S. Ito, M. Myroshnychenko, S. Nigam, M. Shimono, F.-C. Yeh, P. Hottowy, A. M. Litke, and J. M. Beggs. High-degree neurons feed cortical computations. PLOS Computational Biology, 12(5):1–31, 05 2016.

71. J. Touboul. and A. Destexhe. Can power-law scaling and neuronal avalanches arise from stochastic dynamics? PLOS ONE, 5(2):1–14, 02 2010.

72. J. Touboul. and A. Destexhe. Power-law statistics and universal scaling in the absence of criticality. Phys. Rev. E, 95:012413, Jan 2017.

73. E. K. Towlson, P. E. Vertes, G. Yan, Y. L. Chew, D. S. Walker, W. R. Schafer, and A. L. Barabasi. Caenorhabditis elegans and the network control framework-FAQs. Philos Trans R Soc Lond B Biol Sci, 373:1758, 2018.

74. X.-J. Wang, Probabilistic decision making by slow reverberation in cortical circuits. Neuron, 36(5):955–968, 2018/09/04 2002.

75. Y. Wang, H. Markram, P. H. Goodman, T. K. Berger, J. Ma, and P. S. Goldman-Rakic. Heterogeneity in the pyramidal network of the medial prefrontal cortex. Nature Neuroscience, 9:534 EP –, 03 2006.

76. Z. Wang, L. M. Chen, L. Négyessy, R. M. Friedman, A. Mishra, J. C. Gore, and A. W. Roe. The relationship of anatomical and functional connectivity to resting-state connectivity in primate somatosensory cortex. Neuron, 78(6):1116 – 1126, 2013.

77. D. J. Watts and S. H. Strogatz. Collective dynamics of ‘small-world’networks. Nature, 393:440 EP –, 06 1998.

78. L. Wiles, S. Gu, F. Pasqualetti, B. Parvesse, D. Gabrieli, D. S. Bassett, and D. F. Meaney. Autaptic connections shift network excitability and bursting. Sci Rep, 7:44006, 2017.

79. E. Wu-Yan, R. F. Betzel, E. Tang, S. Gu, F. Pasqualetti, and D. S. Bassett. Benchmarking measures of network controllability on canonical graph models. Journal of Nonlinear Science, Mar 2018.

80. G. Yan, P. E. Vértes, E. K. Towlson, Y. L. Chew, D. S. Walker, W. R. Schafer, and A.-L. Barabási. Network control principles predict neuron function in the caenorhabditis elegans connectome. Nature, 550:519 EP –, 10 2017.

81. E. Yeger-Lotem, S. Sattath, N. Kashtan, S. Itzkovitz, R. Milo, R. Y. Pinter, U. Alon, and H. Margalit. Network motifs in integrated cellular networks of transcription-regulation and protein-protein interaction. Proc Natl Acad Sci U S A, 101(16):5934–5939, 2004.

