## Supplemental Information for "Network topology of neural systems supporting avalanche dynamics predicts stimulus propagation and recovery"

**Summary of content.** In this supplementary document we provide various paralipomena relevant to the study's methods, results, and conclusions. In the Supplementary Methods section, we tabulate the parameters used in all network simulations whose results are reported in the main text (see Table 1). These parameters include the number of neurons, the fractional connectivity, the graph model, and the choice of edge weighting scheme. In the Supplementary Results section, we (i) reproduce the results from Figure 3g-i in the main text, but with different sets of edges (see Figure 1), (ii) demonstrate that the finite average controllability is correlated with the mean cascade duration for higher values of $F$ than those reported in the main manuscript (see Figure 2), and (iii) reproduce the results from Figure 4e-j in the main text with networks of higher fractional connectivities (see Figure 3). Finally, in the Supplementary Discussion section, we make note of semantic differences between the terms *avalanche* and *cascade* in the context of the computational neuroscience literature.

### Supplementary Methods

#### Parameters used in network simulations.

Table 1 displays parameters of the network simulations reported in the main text.

| Simulation parameters | | | | |
|---|---|---|---|---|
| Simulation | Number of neurons | Fractional connectivity | Topology | Weighting |
| Figure 1a-e | 10 | 0.2 | WR | uniform |
| Figure 2f | 50, 100, 150, 200, 250, 300 | 0.2 | WR | uniform |
| Figure 2f | 10 | 0.2 | WR | uniform |
| Figure 2 extended | 12 | 0.2 | WR, RG, M4C, WS | uniform |
| Figure 3a | 3 | 0.22 | WR | uniform |
| Figure 3b | 3 | 0.33 | WR | uniform |
| Figure 3c | 10 | 0.45 | WR | uniform |
| Figure 3d-g | 2 | 1.0 | cyclic | sweep |
| Figure 3h-k | 4 | 0.5 | cyclic | sweep |
| Figure 4a-d | 100 | 0.2 | WR | uniform |
| Figure 4e-h | 100 | 0.2 | WR | bimodal Gaussian |
| Figure 5e-j | 100 | 0.2 | WR, RG, M4C, WS | bimodal Gaussian |

**Table 1. Network parameters for all simulations.** The graph topologies are weighted random (WR), random geometric (RG), modular with 4 communities (M4C), and Watts-Strogatz (WS).

### Supplementary Results

**Random redistribution from 4-node cycles to different sets of edges.** In this section, we reproduce the results from Figure 3g-i in the main text, but with different sets of edges (see Figure 1 in this supplementary document). The results provided here are qualitatively similar to those shown in Figure 3 of the main text.

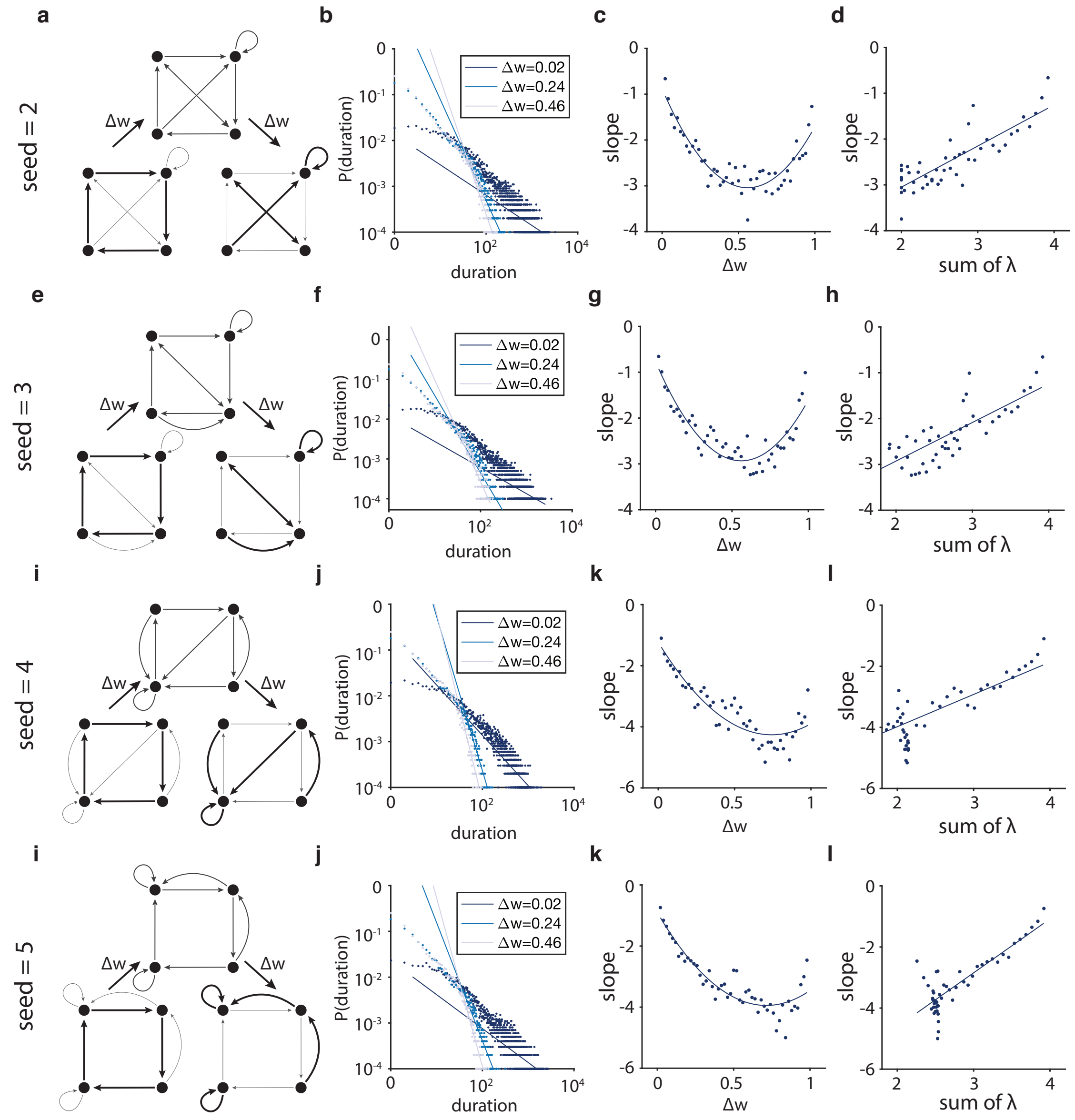


**Figure 1. Strong edge weights in cycles produce long cascades.** Panels **a-d** reproduce the results from Figure 3h-k in the main text with a different set of edges generated by a random seed of 2. The subpanels **a**, **b**, **c**, and **d** here correspond to subpanels h, i, j, and k of Figure 3 in the main text. Panels **e-h** reproduce the results from Figure 3h-k with a different set of edges generated by a random seed of 3. Panels **i-l** rreproduce the results from Figure 3h-k with a different set of edges generated by a random seed of 4. Panels **m-p** reproduce the results from Figure 3h-k with a different set of edges generated by a random seed of 5.

**Finite average controllability for different time periods.** We demonstrate that the finite average controllability is correlated with the mean cascade duration even with higher values of $F$, a parameter reflecting the time period over which the system's impulse response is measured. In the literature [5], average controllability is defined as Trace($W_K$) where $W_K = \sum_{\tau=0}^{\infty} A^\tau B_K B_K^{\mathrm{T}} A^\tau$. Because avalanches are expected to last for a finite number of time steps, we define finite average controllability as the trace of a finite version of the controllability Gramian, $W_K = \sum_{\tau=0}^{\mathrm{F}} A^\tau B_K B_K^{\mathrm{T}} A^\tau$, as discussed more fully in the Methods section of the main text. Here we show that higher values of $F$ only increase the correlation between mean cascade duration from a stimulus and the finite average controllability value of the stimulus (see Figure 2 in this supplementary document). For readers curious about pragmatic concerns, we note that the estimation of finite average controllability becomes much more computationally intensive as $F$ increases.

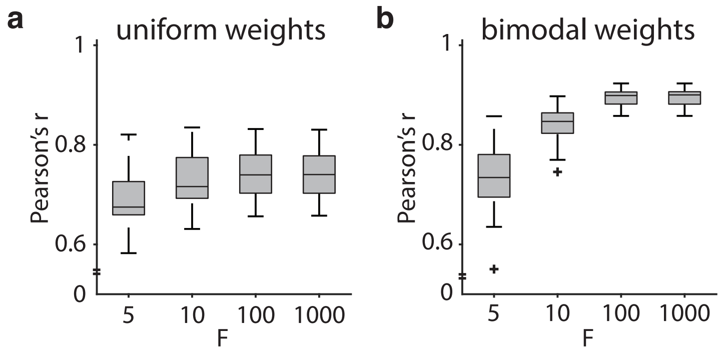


**Figure 2. Finite average controllability, estimated for a wide range of $F$ values, is correlated with mean cascade duration. a,** The Pearson's correlation coefficient, $r$, between the mean duration of cascades from a single-node stimulation and the finite average controllability, for values of $F$ ranging from 5 to 1,000. The networks here have a weighted random network topology with fractional connectivity of 0.2 and a uniform distribution of weights. **b,** The Pearson's correlation coefficient, $r$, for the same measurements as in panel **a** except for a bimodal distribution of weights, as explained in the main text.

**Mutual information decay calculations for networks with different fractional connectivities.** Here we reproduce the results from Figure 4e-j in the main text with networks of higher fractional connectivities of 0.1 and 0.2. The networks with higher fractional connectivities shown here display similarly high correlations between (i) the decay in mutual information between stimuli and network state over time and (ii) the mean duration of cascades (see Figure 3 in this supplementary document).

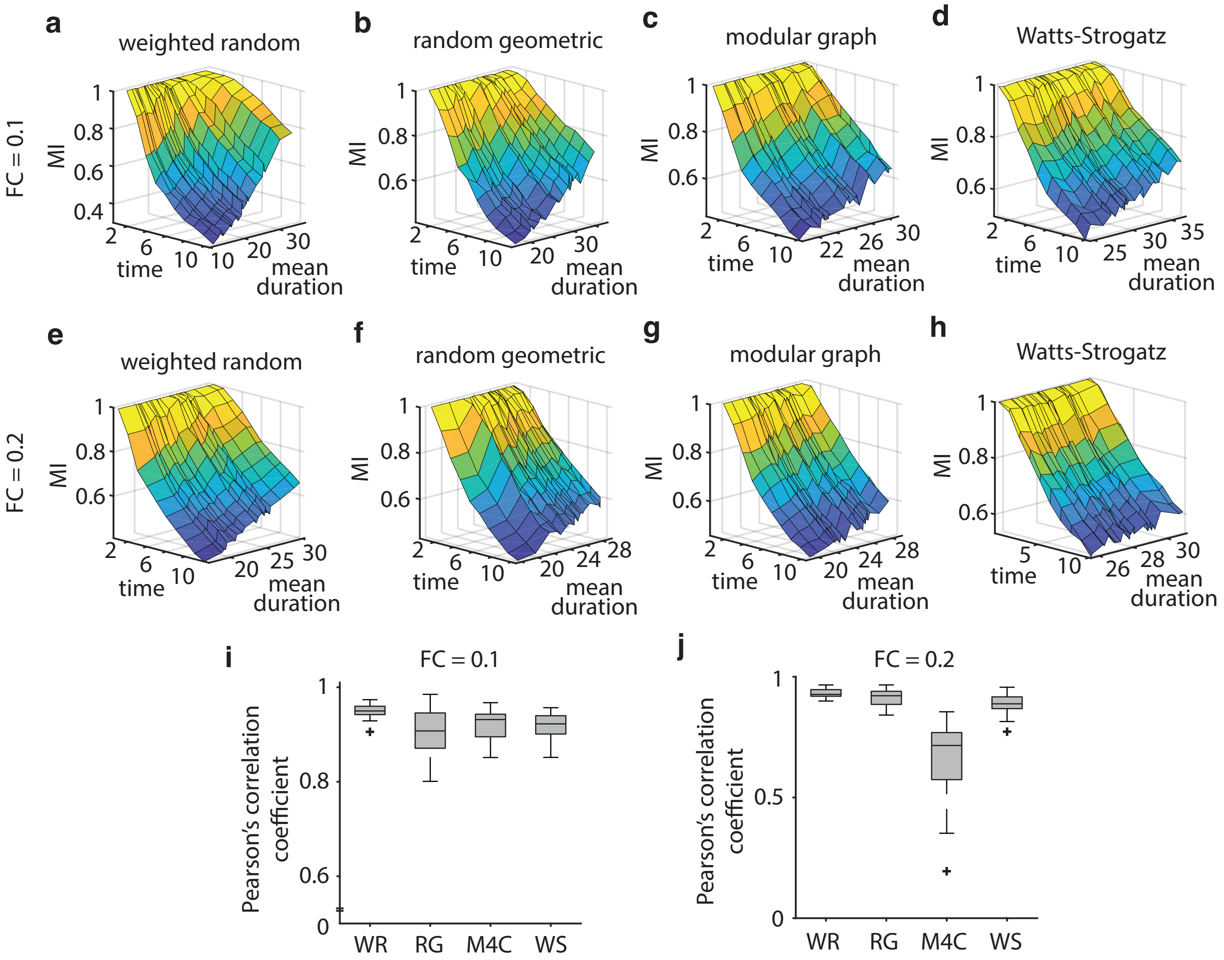


**Figure 3. Longer cascade duration allows stimulus recovery. a-d,** Decay in mutual information (MI) over time. When activity from a stimulus pattern lasts longer, mutual information also persists for longer. Panels show results from four graph types: a weighted random graph, a random geometric graph, a modular graph with 4 communities, and a Watts-Strogatz graph. The networks for these panels have fractional connectivity of 0.1 and a bimodal distribution of weights. **e-h,** The same results are presented here as in panels **a-d** for networks with a fractional connectivity of 0.2. **i,** A boxplot of the Pearson correlation coefficients between the linear slope of decay in mutual information over time and the mean cascade duration. The boxplot shows data from 30 instantiations of each graph type, each network containing 100 nodes and characterized by a fractional connectivity of around 0.1. **j,** The same results are presented here as in panel **i** for networks with fractional connectivity of 0.2.

#### Supplementary Discussion

**Neuronal avalanches versus cascades.** Neuronal avalanches are cascades of spontaneous neuronal activity that follow a power law distribution of sizes and durations that is typical of avalanches and other critical systems [1, 2]. Neuronal cascades, however, do not always display critical behavior. While empirical distributions of cascade size seem to follow power laws, empirical distributions of cascade duration, also referred to as life times, display a wide range of power law exponents from -1.0 to -2.6 [2–4, 6–11]. Moreover, it is clear even without rigorous statistical methods that many empirical distributions of event duration do not follow power laws, but more closely resemble exponential distributions. Such empirical observations beg the question of whether cascades of activity should be called neuronal avalanches. Thus, it seems more appropriate to call cascades of neuronal activity "neuronal cascades," and in the paper, we avoid the assumption of criticality in our simulations and analyses.

### References


1. P. Bak, C. Tang, and K. Wiesenfeld. Self-organized criticality: An explanation of the 1/f noise. *Physical Review Letters*, 59(4):381–384, July 1987.

2. J. M. Beggs and D. Plenz. Neuronal avalanches in neocortical circuits. *Journal of Neuroscience*, 23(35):11167–11177, 2003.

3. T. Bellay, A. Klaus, S. Seshadri, and D. Plenz. Irregular spiking of pyramidal neurons organizes as scale-invariant neuronal avalanches in the awake state. *eLife*, 4:e07224, jul 2015.

4. N. Friedman, S. Ito, B. A. W. Brinkman, M. Shimono, R. E. L. DeVille, K. A. Dahmen, J. M. Beggs, and T. C. Butler. Universal critical dynamics in high resolution neuronal avalanche data. *Phys. Rev. Lett.*, 108:208102, May 2012.

5. S. Gu, F. Pasqualetti, M. Cieslak, Q. K. Telesford, A. B. Yu, A. E. Kahn, J. D. Medaglia, J. M. Vettel, M. B. Miller, S. T. Grafton, and D. S. Bassett. Controllability of structural brain networks. *Nature Communications*, 6:8414 EP –, 10 2015.

6. G. Hahn, T. Petermann, M. N. Havenith, S. Yu, W. Singer, D. Plenz, and D. Nikolić. Neuronal avalanches in spontaneous activity in vivo. *Journal of Neurophysiology*, 104(6):3312–3322, 2010. PMID: 20631221.

7. F. Lombardi, H. J. Herrmann, D. Plenz, and L. De Arcangelis. On the temporal organization of neuronal avalanches. *Frontiers in Systems Neuroscience*, 8:204, 2014.

8. T. Petermann, T. C. Thiagarajan, M. A. Lebedev, M. A. L. Nicolelis, D. R. Chialvo, and D. Plenz. Spontaneous cortical activity in awake monkeys composed of neuronal avalanches. *Proceedings of the National Academy of Sciences*, 106(37):15921–15926, 2009.

9. S.-S. Poil, R. Hardstone, H. D. Mansvelder, and K. Linkenkaer-Hansen. Critical-state dynamics of avalanches and oscillations jointly emerge from balanced excitation/inhibition in neuronal networks. *Journal of Neuroscience*, 32(29):9817–9823, 2012.

10. A. Ponce-Alvarez, A. Jouary, M. Privat, G. Deco, and G. Sumbre. Whole-brain neuronal activity displays crackling noise dynamics. *Neuron*, 11 2018.

11. W. L. Shew, W. P. Clawson, J. Pobst, Y. Karimipanah, N. C. Wright, and R. Wessel. Adaptation to sensory input tunes visual cortex to criticality. *Nature Physics*, 11:659 EP –, 06 2015.
